# TRPA1 block protects against loss of white matter function during ischaemia in the mouse optic nerve

**DOI:** 10.1101/2021.05.25.445536

**Authors:** Wendy Lajoso, Grace Flower, Vincenzo Giacco, Anjuli Kaul, Circe La Mache, Andra Brăban, Angela Roxas, Nicola Hamilton

**Affiliations:** Wolfson Centre for Age-Related Diseases, Institute of Psychiatry, Psychology and Neuroscience, Guy’s Campus, King’s College London, SE1 1UL, UK

## Abstract

Oligodendrocytes produce myelin which provides insulation to axons and speeds up neuronal transmission. In ischaemic conditions myelin is damaged, resulting in mental and physical disabilities. Therefore, it is important to understand how the function of oligodendrocytes and myelin is affected by ischaemia. Recent evidence suggests that oligodendrocyte damage during ischaemia is mediated by TRPA1, whose activation raises intracellular Ca^2+^ concentrations and damages compact myelin. Here, we show that TRPA1 is constitutively active in oligodendrocytes and the optic nerve, as the specific TRPA1 antagonist, A-967079, decreases basal oligodendrocyte Ca^2+^ concentrations and increases the size of the compound action potential. Conversely, TRPA1 agonists reduce the size of the optic nerve compound action potential, and this effect is significantly reduced by the TRPA1 antagonist. These results indicate that glial TRPA1 regulates neuronal excitability in the white matter under physiological as well as pathological conditions. Importantly, we find that inhibition of TRPA1 prevents loss of compound action potentials during oxygen and glucose deprivation (OGD) and improves the recovery. TRPA1 block was effective when applied before, during or after OGD, indicating that the damage is occurring during ischaemia and during the recovery, but importantly, that therapeutic intervention is possible after the ischaemic insult. These results indicate that TRPA1 has an important role in the brain, and that its block may be effective in treating oligodendrocyte loss and damage in many white matter diseases.

## Introduction

Oligodendrocytes (OLs), wrap fatty myelin sheaths around axons to decrease the capacitance across the axonal membrane and increase the action potential speed. Myelin loss in diseases such as periventricular leukomalacia, leukodystrophies, multiple sclerosis, and stroke, leads to failure of neuronal transmission and thus mental and physical impairment. We have recently shown that cerebellar oligodendrocytes express transient receptor potential ankyrin-1 (TRPA1), whose activation during ischaemia causes excessive Ca^2+^ influx and myelin damage (Hamilton et al., 2016).

TRPA1 is one of a large family of tetrameric non-selective cation channels that are widely expressed in the grey and white matter of the CNS and are increasingly considered as potential therapeutic targets in brain disorders (Cornillot et al., 2019). TRPA1 is sensitive to environmental irritants and endogenous electrophilic compounds that are formed during oxidative stress (Andersson et al., 2008), which have been shown to evoke pain, cold, itch and inflammation (Bautista et al., 2013). Like many TRP channels, TRPA1 is highly permeable to Na^+^ and Ca^2+^, which in turn can deregulate cell function and cause apoptosis. In cerebellar OLs, TRPA1 was shown to be activated during ischaemia via acidification of the cytosol (Hamilton et al., 2016). However, until now we did not know whether OLs in other areas of the brain express functional TRPA1 and whether TPRA1 block is protective against loss of white matter function in ischaemia.

Here, we show by patch-clamping corpus callosum OLs, that they also respond to TRPA1 agonists by raising their intracellular Ca^2+^ concentrations ([Ca^2+^]_i_), and that TRPA1 inhibition with A967079 (2μM) reduces the resting [Ca^2+^]_i_. Furthermore, the TRPA1 antagonist also increases the amplitude of optic nerve compound action potential recordings under normal physiological conditions and prevents a large proportion of the OGD induced loss of the action potential. This shows that TRPA1 may play a role in regulating neuronal excitability and TRPA1 inhibition may be a possible treatment for white matter damage in diseases where acute or chronic ischaemia has been implicated.

## Methods

### Animals

C57BL/6J mice of either sex were killed via schedule 1 (cervical dislocation) in accordance with the guidelines of the UK Animals (Scientific Procedures) Act 1986 and subsequent amendments. TRPA1 knockout (KO) mice originated from Kelvin Kwan, Rutgers University and are now available here (https://www.jax.org/strain/008649). They were kindly provided to us by John Wood, Stuart Bevan and David Andersson, and then bred in house from heterozygous pairs to provide KOs and littermate wildtype (WT) controls. These mice were created by putting loxP sites on either side of exons 22-24 (encoding the S5/S6 transmembrane domains), which produces a dysfunctional TRPA1 channel in these mice.

### Brain slice and optic nerve preparation

Coronal brain slices (225 μm thick) were prepared from the brains of P12-P17 mice in icecold solution containing (mM) 124 NaCl, 26 NaHCO_3_, 1 NaH_2_PO_4_, 2.5 KCl, 2 MgCl_2_, 2 CaCl_2_, 10 glucose, bubbled with 95% O_2_/5% CO2, pH 7.4, as well as 1 mM Na-kynurenate to block glutamate receptors. Optic nerves were dissected from P42-P70 mice. Brain slices and optic nerves were then incubated at room temperature (21–24 °C) in the above solution until used in experiments.

### Cell identification and electrophysiology

Oligodendrocytes were identified by their location and morphology. All cells were wholecell clamped with pipettes with a series resistance of 8-35 MΩ. Electrode junction potentials were compensated and cells were voltage-clamped at −74 mV.

### External solutions

Slices and optic nerves were superfused with bicarbonate-buffered solution containing (mM) 124 NaCl, 2.5 KCl, 26 NaHCO_3_, 1 NaH_2_PO_4_, 2 CaCl_2_, 1 MgCl_2_, 10 glucose, pH 7.4, bubbled with 95% O_2_ and 5% CO_2_. Applications of TRPA1 channel agonists and antagonists to brain slices were at room temperature. All compound action potential recording and OGD experiments were performed between 33 and 36 °C. Control and drug conditions were interleaved and the experimenter was blinded to the solution contents. To simulate ischaemia we replaced external O2 with N2, and external glucose with 7 mM sucrose. The flow rate was approximately 4 mL per minute into a 1.5 mL bath, giving a turnover rate of under 25 seconds.

### Intracellular solutions

Cells were whole-cell clamped with electrodes containing K-gluconate-based solution, comprising (mM) 130 K-gluconate, 2 NaCl, 0.5 CaCl_2_, 10 HEPES, 10 BAPTA, 2 NaATP, 0.5 Na_2_GTP, 2 MgCl, and 0.05 Alexa Fluor 594 pH set to 7.15 with K-OH (all from Sigma). For Ca^2+^ imaging experiments, BAPTA was decreased to 0.01 mM and replaced with 10 mM phosphocreatine, CaCl_2_ was reduced to 10 μM, and 200 μM Fluo-4 or Fluo-8 was added to allow ratiometric imaging with the above Alexa Fluor.

### Single cell ion imaging

Fluo-8 and Alexa Fluor 594 were used in the internal solution to measure [Ca^2+^]_i_ changes ratiometrically during experiments. Fluo-8 and Alexa Fluor 594 fluorescence were excited sequentially using a monochromator every 3 seconds at 488 ± 10nm and 585 ± 10nm, and emission was collected using a triband filter cube (DAPI /FITC/Texas Red, 69002, Chroma). The mean ratio of intensities (excited at 488 nm/excited at 585 nm) before applying the TRPA1 agonist was 1.010 (n = 28).

### Drug application

Stock solutions of the following drugs were made up: Carvacrol (Sigma) was diluted in 100% ethanol, HC 030031 (Tocris), A-967079 (Boc Sciences Laboratory), flufenamic acid (Sigma), polygodial (Tocris) and AITC (Sigma) were made up in 100% DMSO. For CAP experiments, DMSO and ethanol were also added to control solutions at the same concentrations and did not evoke any changes in the CAP amplitude at the concentrations used (≤0.1%). For the patch-clamping experiments, vehicle controls have been tested on oligodendrocytes and do not evoke any changes at the concentrations used. Stocks were kept at −20°C apart from carvacrol and AITC, which were made up fresh on each day of use. To minimise evaporation of carvacrol, lids were kept on until the solutions were used. The control and test experiments were interleaved, and for all CAP recordings, the experimenter was blinded to the contents of the solutions.

### Compound action potential recording

Optic nerves were isolated from P42-P70 mice and compound action potentials recorded using suction electrodes (Bolton and Butt, 2005). The optic nerve was stimulated using a suction electrode filled with extracellular solution, using 0.2 ms, 50-100 V pulses, at 1 Hz to produce as supramaximal a response as possible. Supramaximal stimulation (40% over the voltage producing the maximal amplitude of CAP) was constrained by a maximum stimulation voltage of 100 V. The compound action potential was recorded as a voltage using an Axon Axoclamp 2B amplifier, with the recording suction electrodes placed as far away from the stimulating electrode as possible to obtain a CAP waveform with three peaks, which denote heavily myelinated axons (peak 1), normally myelinated axons (peak 2) and unmyelinated axons (peak 3; (Stys et al., 1991). At the end of each experiment, TTX (500 nM) was applied to obtain the stimulus artefact in the absence of action potentials, which was subtracted from all the records obtained previously. The prestimulus baseline of the resulting traces was subtracted, then they were rectified (squared, then square rooted), and either the area of the compound action potential was calculated or the amplitude of the second peak was measured (see results).

### Statistics

Data are meam ± s.e.m. P values are from ANOVA tests (for normally distributed data) and Mann-Whitney U or Kruskal-Wallis tests (for non-normally distributed data). Normally distributed data were tested for equal variance (p<0.05, unpaired F-test) and paired t-tests were adjusted accordingly. P values quoted in the text are from ANOVA tests unless otherwise stated. For multiple comparisons, P values were corrected using the Holm-Bonferroni or Tukey’s test. Small sample sizes were often able to achieve statistical significance, so in those instances power analysis of sample size was also measured for soma data and determined to be greater than 0.75. Data normality was assessed using Shapiro-Wilk, Anderson-Darling, D’agostino, and Kolmogorov-Smirnov tests. All statistical analysis was conducted using OriginLab or GraphPad Prism software.

## Results

### Oligodendrocytes express tonically active TRPA1

The corpus callosum (CC) is a white matter tract spanning across the two hemispheres that is often thinned or demyelinated as a result of local hypoperfusion occurring in periventricular leukomalacia or stroke victims (Yang et al., 2014). Therefore, our first aim was to determine whether OLs in the CC also express functional TRPA1. To do this we whole cell patch-clamped OLs identified at first by their morphology (light oval somata laid in lines within parallel axons) and then by their dye-filled morphology (n = 28, Fig. 1a). Using the patch-pipette, the OLs were dye filled with the Ca^2+^ sensitive dye Fluo-8 (200 μM). This method ensures that the measured [Ca^2+^]_i_ changes are in OLs only. Like previously reported in cerebellar OLs (Hamilton et al., 2016), the TRPA1 agonists flufenamic acid (FFA, 100 μM) and carvacrol (2mM) evoke a [Ca^2+^]_i_ increase in OLs in the corpus callosum (P12-17; Fig. 1b,d). However, while the patch pipette was present, the TRPA1 agonists AITC (500 μM), polygodial (100 μM) did not. When we repeated the experiments with the patch-pipette removed, a short while after loading the cell with the Ca^2+^ indicators, we found that AITC and polygodial were then able to activate TRPA1 (Fig. 1c,d). This lack of response to this specific group of TRPA1 agonists during patch-clamping techniques has been found previously (Kim and Cavanaugh, 2007) and suggests that a necessary intracellular component may be washed out or chelated in these instances that modifies the availability of the agonist binding site. These agonists covalently bind to cysteine residues found on the N-terminal. Importantly, application of the TRPA1 antagonists A967079 (20 μM) or HC-030031 (100 μM) decreases the resting [Ca^2+^]_i_ in both corpus callosum and cerebellar OLs, as it does in astrocytes (Shigetomi et al., 2012), suggesting that TRPA1 in OLs is tonically active (Fig. 1e) and regulates normal cell functions.

**Figure 1.**
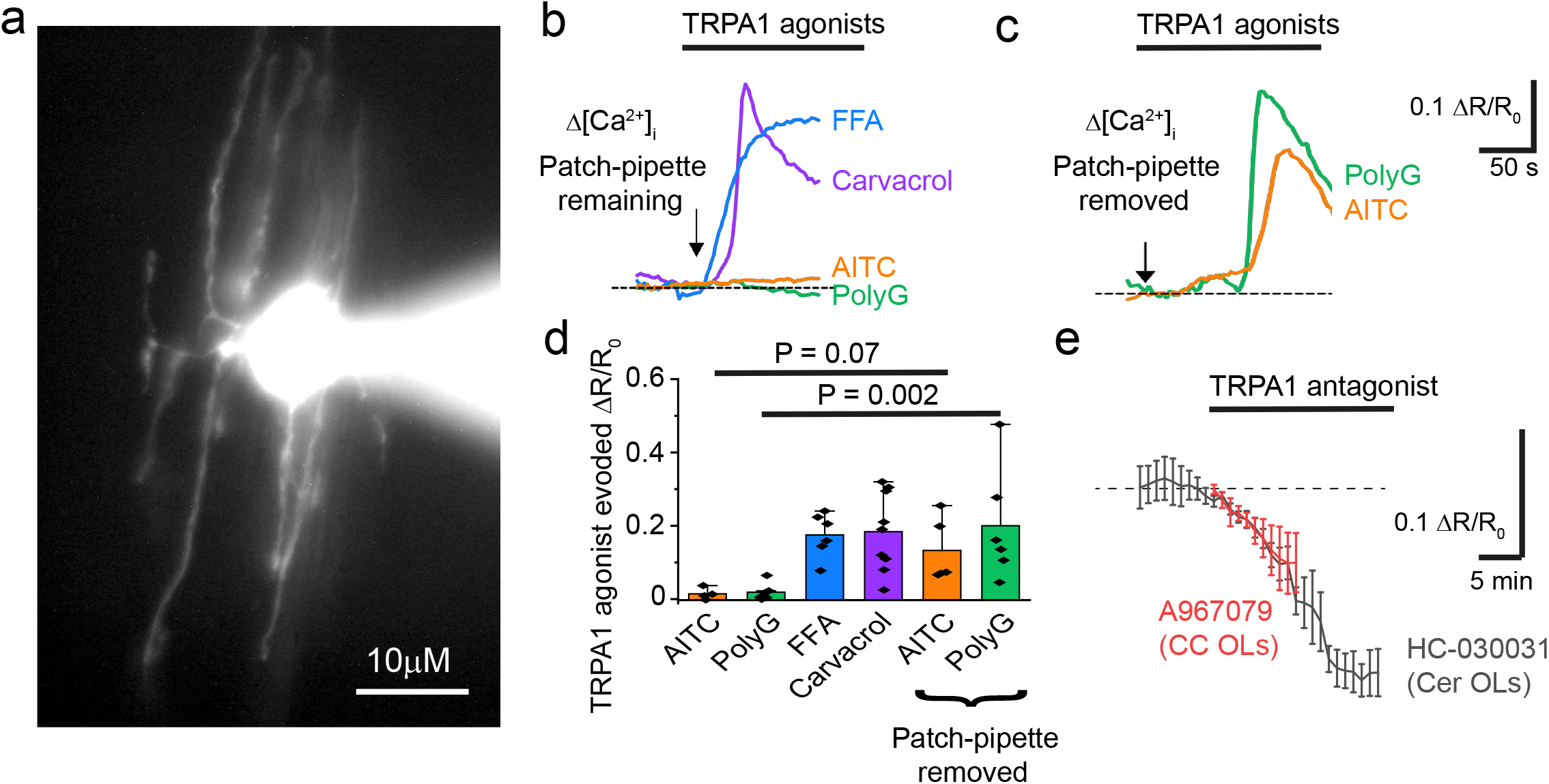
Corpus callosum oligodendrocytes express TRPA1. **(a)** Whole-cell patch clamped oligodendrocyte (OL) with Alexa dye in soma and processes. **(b)** Δ [Ca^2+^]_i_ to TRPA1 agonists with the patch-pipette remaining: FFA 100 μM (n=6); carvacrol 2mM (n = 8); 500 μM AITC (n = 4); polygodial 100 μM (n = 6). **(c)** Δ [Ca^2+^]_i_ to TRPA1 agonists with patch-pipette removed: AITC 500 μM (n = 5); polygodial 100 μM (n = 6). **(d)** TRPA1 agonists evoke in increase in Δ [Ca^2+^]_i_ (P values compare with and without patch pipette, one-way ANOVA test with Holm-Bonferroni correction for multiple comparisons). **(e)** A decrease in [Ca^2+^]_i_ occurs in in the presence of TRPA1 antagonists: A967079 20 μM in corpus callosum (CC) OLs and HC-030031 100 μM in cerebellar (Cer) OLs.

### Tonic TRPA1 activation regulates the compound action potential amplitude in the optic nerve

White matter Ca^2+^ concentrations, and receptors that allow Ca^2+^ flux have been suggested to regulate action potential propagation in disease (Stys et al., 1990, 1991; Micu et al., 2006; Bakiri et al., 2008; Baltan, 2016). As TRPA1 regulates intracellular Ca^2+^ concentrations in oligodendrocytes, we hypothesised that tonic TRPA1 activity may have an effect on oligodendrocyte function in maintaining fast neuronal action potential propagation. We used the optic nerve to measure compound action potential amplitude as described previously (Bolton and Butt, 2005; Bakiri et al., 2008; Fig.2a) because it is a heavily myelinated, easily accessible CNS white matter tract (Stys et al., 1991). Interestingly, we discovered a modest regulation of CAP amplitude by tonic TRPA1 activity in the optic nerve as TRPA1 inhibition with A-967079 (20 μM) increased the amplitude by 12 ± 4% within 10 minutes (n = 9; Fig. 2b,c). Conversely, the TRPA1 agonists AITC (500 μM; n = 8; Fig. 2d,e), and polygodial (10 or 100 μM; n = 6-7; Fig. 2f) reduced the size of the compound action potential amplitude (n = 5 - 14; Fig. 2g). The TRPA1 agonist-induced decrease in compound action potential amplitude was reduced in the presence of A-967079, which was preincubated for over 15 minutes to allow for the initial increase in CAP amplitude before application of the agonists (n = 5 - 14; Fig. 2g). As optic nerves do not include neuronal somata, these results indicate glial TRPA1 regulates retinal ganglion cell compound action potential propagation in physiological conditions. Whether this is due to OL or astrocyte TRPA1 is yet to be determined.

**Figure 2.**
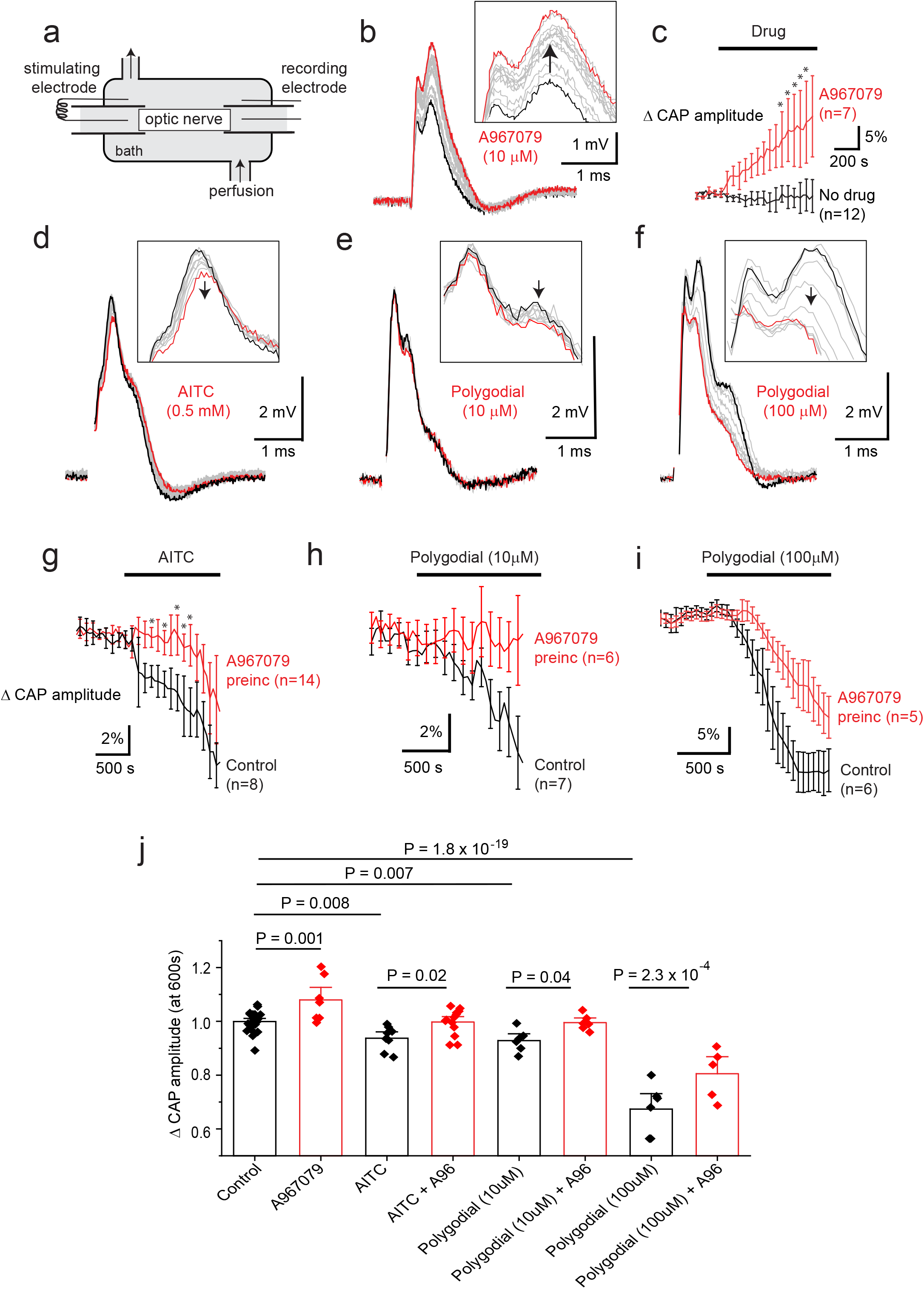
TRPA1 agonist and antagonist effects on optic nerve compound action potential amplitude. **(a)** Compound action potentials (CAP) were measured in young adult optic nerves (P42 – P70). **(b)** A967079 (A96, 10 μM) block of TRPA1 increases the CAP amplitude (black trace at start, with time this increases (grey traces) and the red trace is the final trace). **(c)** Mean CAP amplitude increases after application of A967079 (10 μM; n = 7) compared to vehicle (DMSO; n = 12) and decreases after TRPA1 agonist application: AITC (500 μM, d, g, j, n =8); and polygodial (10 μM, **e h, j**, n = 7; and 100 μM, **f, i, j**, n = 6). TRPA1 – evoked decreases can be significantly inhibited by over 15 minutes of preincubation with A967079 (20 μM, **g-j**). **(j)** Mean changes in CAP amplitude after application of agonists and antagonists for 10 minutes. P values are from multiple comparisons using a one-way ANOVA with Holm-Bonferroni correction.

### TRPA1 block protects against loss of the compound action potential during OGD

In the past, CAP recordings have demonstrated that simulated ischaemia results in a reduction in action potential propagation with only a partial recovery (Bastian et al., 2020; Dong and Hare, 2005; Stys et al., 1990; Hamner et al., 2015; Bakiri et al., 2008). This reduction in CAP amplitude is thought to be mediated by Ca^2+^ influx through different channels, therefore TRPA1 inhibition may protect against loss of compound action potentials during ischaemia. To simulate ischaemia we removed O2 and glucose for 30 minutes, and continually recorded the CAP during a recovery period of 30 minutes. The area under the curve of the CAP was measured throughout. To understand the time at which TRPA1 activation may cause the most damage, we applied A-967079 at different times during the experiment. Firstly, we pre-incubated A-967079 (20 μM) and administered it throughout the experiment and found it significantly reduced the loss of the CAP during ischaemia (Fig. 4a, d; P = 0.02), and improved the recovery from 61 ± 3.7 % to 89 ± 7.4% (Fig. 4a,e; P = 7.5 × 10^−5^). As therapeutics are more often than not given either during or after the event rather than prophylactically, we applied A-967079 during either the ischaemic insult or only the recovery period (Fig. 4b-e). A-967079 application during the ischaemic insult or the recovery period alone resulted in an improved CAP recovery (77 ± 8 %; P = 0.02 and 73.7 ± 4 %; P = 0.05 respectively) suggesting that the TRPA1 mediated damage is occurring during both the ischaemic and recovery periods. These results suggest that TRPA1 activation during ischaemia is a major cause of white matter damage and that TRPA1 antagonists may be effective in treating white matter in the many diseases involving ischaemia.

**Figure 3.**
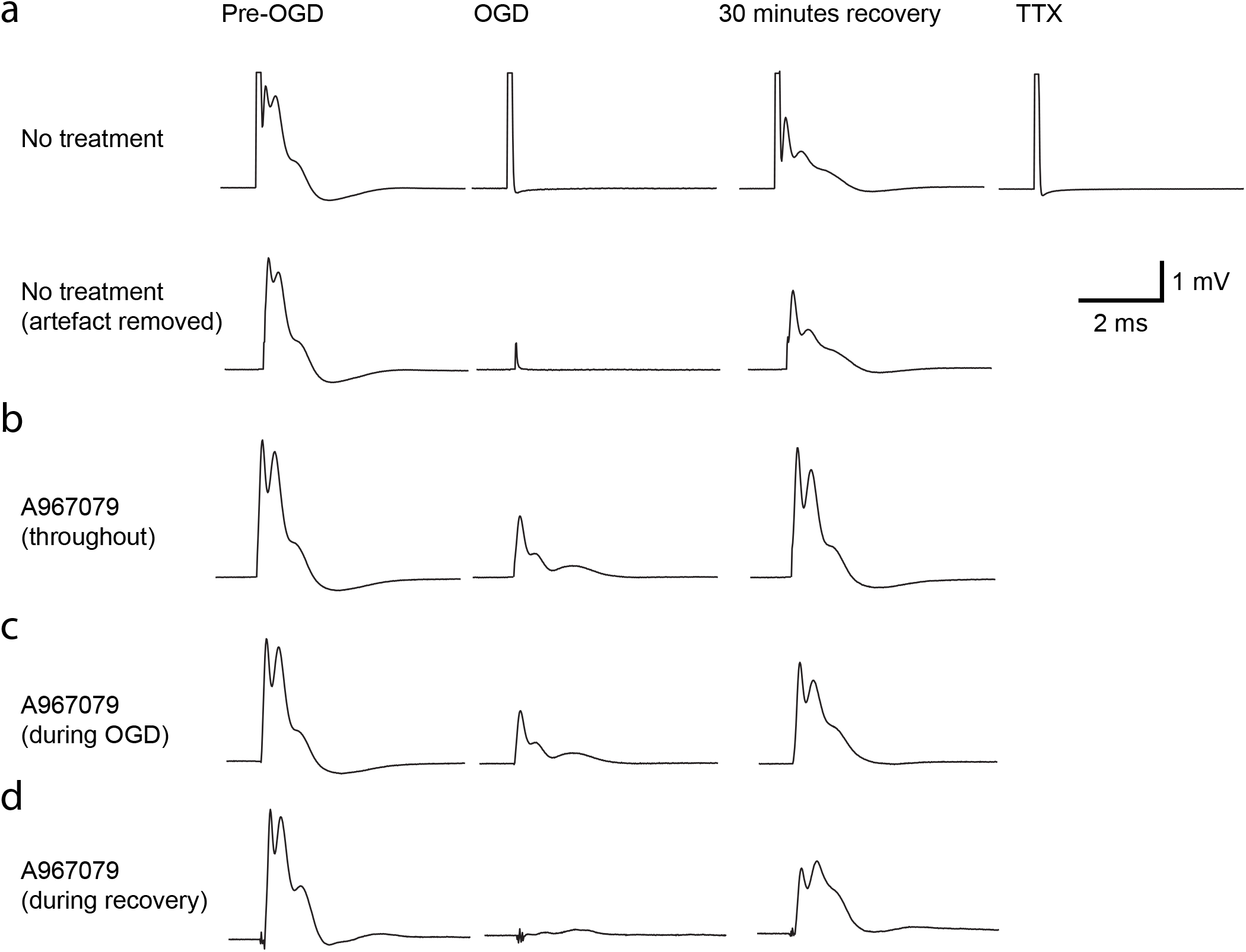
Ischaemia-evoked changes in optic nerve compound action potential (CAP). **(a)** Top row, traces from the left are: CAP in control solution, after 30 minutes ischemia, after 30 minutes recovery to control solution, after application of 500nM TTX to record the stimulus artefact. Bottom row shows the same traces after the stimulus artefact has been removed from the traces. **(b-d)** Artefact subtracted responses with A967079 (20 μM) preincubated and added throughout the experiment **(b)** during ischaemia only **(c)** or during the recovery only **(d)**.

**Figure 4.**
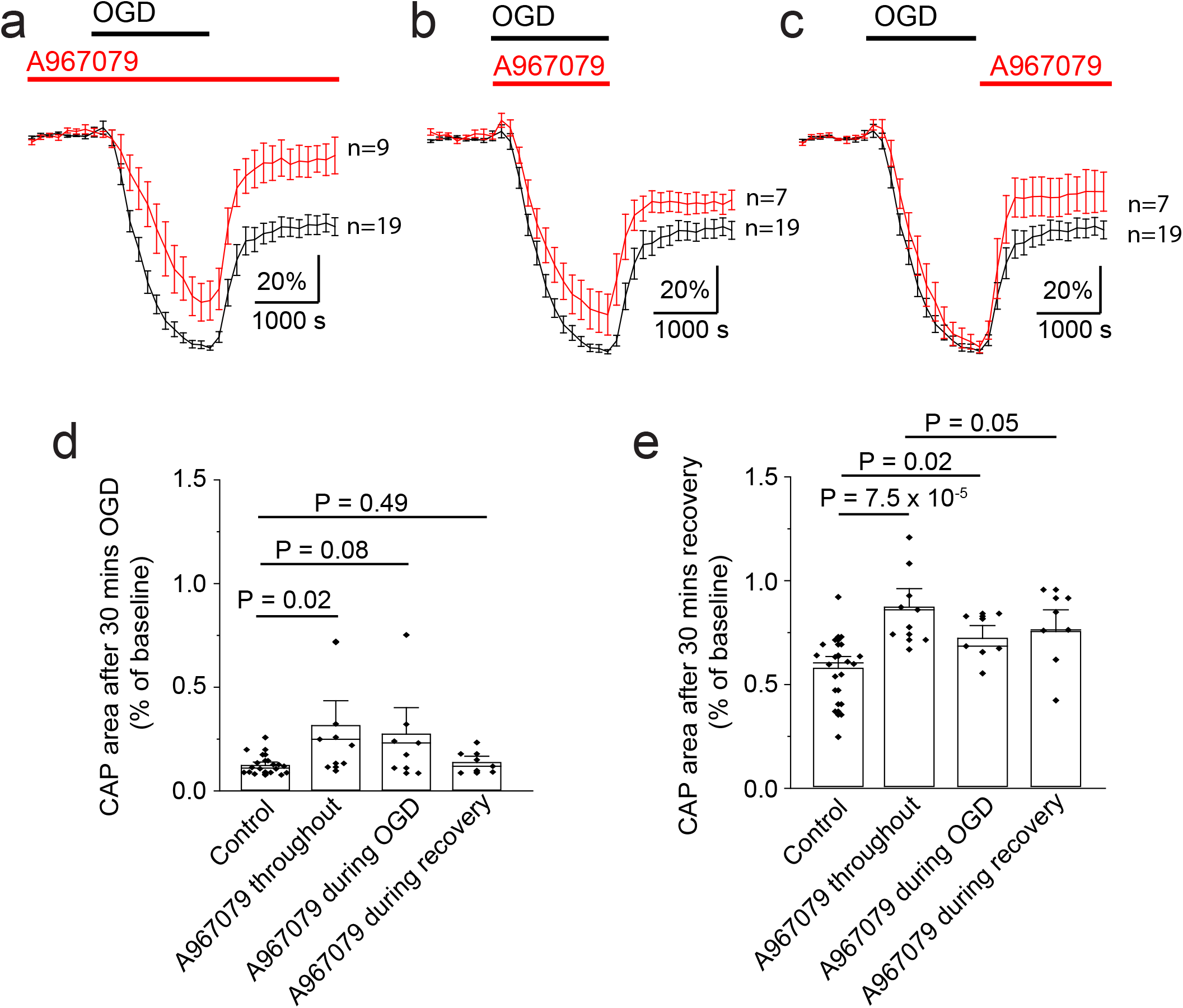
Ischaemia-evoked changes in optic nerve compound action potential (CAP) are reduced by TRPA1 block with A967079. **(a)** Normalised area of CAP before, during and after ischaemia, showing how preincubation of the TRPA1 antagonist (A967079, 20 μM) can prevent the loss of the CAP during ischaemia and improve the recovery (n= 9). Application of A967079 during ischaemia **(b)** or during recovery **(c)** is also protective. **(d)** Normalised CAP area after 30 minutes in OGD or after 30 minutes recover **(e).** P values in **(d)** are from Mann Whitney tests with Holm-Bonferroni correction for multiple comparisons. P values in (e) are from a one-way ANOVA with Holm-Bonferroni correction for multiple comparisons.

## Discussion

The presence of TRPA1 in glial cells is a new concept, but the evidence for functional TRPA1 in both astrocytes (Shigetomi et al., 2012; Shigetomi et al., 2013) and OLs is growing (Hamilton et al., 2016; Marques et al., 2016; Lee et al., 2017; Bölcskei et al., 2018 reviewed in Cornillot et al., 2019). In astrocytes and OLs, TRPA1 activation and inhibition raises and decreases basal intracellular Ca^2+^ (and Mg^2+^) concentrations, respectively (Shigetomi et al., 2013; Hamilton et al., 2016; Fig. 1e). Astrocyte TRPA1 regulates neuronal function (Shigetomi et al., 2012; Shigetomi et al., 2013) and TRPA1 knockout modifies the expression of myelinating proteins (Lee et al., 2017). Here, we add to this evidence by showing that TRPA1 expression in OLs appears to be conserved in other CNS brain areas, and that activation of TRPA1 in corpus callosum OLs generates a substantial Ca^2+^ influx. Despite this, we do not find a large TRPA1-mediated nonspecific cation current (that would reverse at −10 to 0mV) as you would expect when TRPA1 is activated in large numbers on the cells (Ye et al., 2018). Instead, we see a TRPA1-mediated decrease in potassium conductance indicating an inhibition of potassium channels. This has been described in Hamilton et al., 2016. This suggests that the levels of TRPA1 in OLs are low, which may be why its expression is not picked up in bulk brain studies (Cahoy et al., 2008), but is in single cell sequencing (Marques et al., 2016) and *in situ* hybridisation experiments (Hamilton et al., 2016). Interestingly, TRPA1 was not found in mouse optic nerves during qPCR experiments (Papanikolaou et al., 2017). This discrepancy is hard to explain as the mouse ages and strain are similar to those used here (C57BL6; P7, P30-40 vs P12-18 and P42-70 here). Nonetheless, our evidence clearly points out to functional expression of TRPA1 in the mouse optic nerve, and we are providing the first evidence for a role for glial TRPA1 in regulating neuronal transmission through the white matter. Although the focus of this paper has been the expression of TRPA1 in OLs, the effect of TRPA1 on astrocytes and possibly OPCs may be equally important.

How TRPA1 activity decreases the size of the compound action potential under physiological conditions has not been completely elucidated. We know that removing extracellular Ca^2+^ protects against loss of CAPs during ischaemia (Stys et al., 1990), suggesting that influx of Ca^2+^ through TRPA1 may diminish the CAP. However, we also know that TRPA1 decreases oligodendrocyte potassium permeability (Giacco et al., 2021 unpublished), which would in turn decrease the CAP amplitude by decreasing glial cell potassium syphoning away from the periaxonal space (Djukic et al., 2007; Larson et al., 2018). How the CAP is affected by inhibited OL potassium conductance is dependent on the amount that this changes the perinodal potassium concentration. Changes of a few mM can depolarise axons and lead to an increased excitability, but large changes (> 10 mM) can lead to conduction block (Kofuji and Newman, 2004; Rash, 2010). In our model, it appears that tonic TRPA1 activation in normal physiological conditions is already limiting axon excitability, and increased TRPA1 activation with exogenously applied agonists builds upon that phenomenon. Therefore, it appears that either perinodal potassium concentrations are already high enough to prevent action propagation in some axons, or that the OL depolarisation or Ca^2+^ influx through TRPA1 is decreasing the excitability of axons in another way. In support of the former, one action potential is thought to increase the perinodal K^+^ concentration by 1mM (Ransom et al., 2000), and this may increase quickly with successive action potentials if potassium syphoning is hindered.

As mentioned above, increased extracellular Ca^2+^ concentrations are correlated with the amount of white matter damage occurring during ischaemia (Stys et al., 1990; Salter and Fern, 2005; Micu et al., 2006), and a large proportion of the Ca^2+^-mediated damage has long been thought to be due to glutamate excitotoxicity driving Ca^2+^ influx through OL AMPA/KA (Tekkök and Goldberg, 2001; Follett et al., 2000) and NMDA (Salter and Fern, 2005; Káradóttir et al., 2005; Micu et al., 2006; Bakiri et al., 2008) receptors. However, our recent work indicates that the majority of the Ca^2+^ influx into OLs during ischaemia is through TRPA1, which becomes activated when OLs acidify as a result of raised extracellular potassium concentrations occurring during ischaemia (Hamilton et al., 2016). Optic nerves subjected to ischaemia for one hour have TRPA1-mediated myelin damage shown with electron microscopy (Hamilton et al., 2016). In line with that, we find here that TRPA1 inhibition is beneficial at protecting against white matter damage when applied during, after or throughout the ischaemic insult, suggesting that TRPA1 block may be used as a potential prophylactic therapy, or one to improve recovery after a stroke. However, the CAP area never returned to its pre-ischaemic level. This residual loss of the CAP may be due to activation of glutamate receptors, or due to underlying damage to the axons (McCarran and Goldberg, 2007), rather than OLs.

At present, there are a large number of patents for the use of TRPA1 antagonists in human pathologies. A-967079 was chosen here because it is commercially available, highly selective, more readily dissolved in aqueous solution than HC-030031 and can penetrate the nervous system at therapeutic concentrations when administered peripherally, by oral gavage and intraperitoneal injection (Chen et al., 2011). This opens up possibilities for testing the effects of A-967079 *in vivo* in stroke models. However, initial evidence suggests that TRPA1 activation on capillaries may induce vasodilation of arterioles and protect against widespread damage during stroke (Earley, 2012; Pires and Earley, 2018). These results were determined using a conditional endothelial cell TRPA1 knockout (Sullivan et al., 2015). Nonetheless, the evidence of TRPA1 mediated pathology within the parenchyma is growing substantially and indicates that targeted TRPA1 knockout or block within the parenchyma will have major benefits and needs investigating further.

## Acknowledgements

Supported by a European Leukodystrophies Grant (ELA2017-015I4), Medical Research Council New Investigator Grant (MR/S003045/1) and King’s College London. We thank Stuart Bevan and David Andersson for knock-out mice and David Attwell for comments on the manuscript.

